# Previewing a face in the periphery reduces the fN170: Combined EEG and eye-tracking suggests two stages of trans-saccadic predictive processes

**DOI:** 10.1101/468900

**Authors:** Christoph Huber-Huber, Antimo Buonocore, Clayton Hickey, David Melcher

## Abstract

The world appears stable despite saccadic eye-movements. One possible explanation for this phenomenon is that the visual system predicts upcoming input across saccadic eye-movements, based on peripheral preview of the saccadic target. We tested this idea using concurrent electroencephalography (EEG) and eye-tracking. Participants made cued saccades to peripheral upright or inverted face stimuli that could change (invalid preview) or keep their orientation (valid preview) across the saccade. Experiment 1 demonstrated better discrimination performance and a reduced fixation-locked N170 (fN170) with valid than with invalid preview demonstrating integration of pre- and post-saccadic information. Moreover, the early fixation-locked EEG showed a preview face inversion effect suggesting that we perceive pre-saccadic input up to about 170 ms post fixation-onset, at least for face orientation. Experiment 2 replicated Experiment 1 and manipulated the proportion of valid and invalid trials (mostly valid versus mostly invalid, 66.6% to 33.3%) to test whether the preview effect reflected active expectations. A whole-scalp Bayes factor analysis provided evidence for no influence of proportion on the fN170 preview effect. Instead, before the saccade the preview face orientation effect declined earlier in the mostly invalid than in the mostly valid block suggesting some form of pre-saccadic expectations. We conclude that visual stability is achieved by two trans-saccadic integration processes: pre-saccadic prediction, reflected in the pre-saccadic proportion modulation, and early post-saccadic change-detection reflected in the fN170 preview effect.

## 1. Introduction

Visual perception appears surprisingly stable despite being interrupted by saccadic eye movements about three times per second. One source of visual stability may be the integration of pre- and post-saccadic visual information (Helmholtz, 1867; Melcher, 2011; Wurtz, 2008). Recent gaze-contingent experimental designs have revealed that orientation (Ganmor et al., 2015; Wolf and Schütz, 2015; Zimmermann et al., 2017), object size (Valsecchi and Gegenfurtner, 2016), visual motion (Fabius et al., 2016), and whole-object information (Castelhano and Pereira, 2017; Schut et al., 2016) are integrated across saccades in a statistically optimal fashion taking into account the relative reliability of pre-saccadic and post-saccadic input (Ganmor et al., 2015; Herwig, 2015; Wolf and Schütz, 2015). Nonetheless, the time-course of trans-saccadic perception and, in particular, the content of perception immediately after fixation-onset remain controversial (for review, Melcher and Morrone, 2015)

Here, we investigated the time-course of trans-saccadic perception with combined EEG and eye-tracking (Huber-Huber et al., 2016; Kovalenko and Busch, 2016). Using a similar methodology, reading research has discovered a *preview positivity* in the fixation-locked EEG starting at around 200 ms in which the evoked response is more positive for valid than for invalid preview (Dimigen et al., 2012; Kornrumpf et al., 2016; Li et al., 2015), suggesting that pre- and post-saccadic information are compared and integrated by around 200 ms. We investigated whether the preview positivity known from reading research is also elicited by non-word stimuli, namely by faces. One advantage of using face stimuli is that the time course of face processing is well known (e.g. Bentin et al., 1996). In Experiment 1, participants made saccades to peripheral face stimuli. During the saccade, the face orientation (upright, inverted) could change (invalid preview) or remain the same (valid preview). If the reading preview positivity reflected a general trans-saccadic integration mechanism, a similar change in the fixation-locked component should be elicited by preview of the target face. However, we hypothesized that faces might show a different preview effect than words, possibly in the N170 component. Repeated presentation of faces has been shown to reduce the N170 component (Caharel et al., 2009; Ewbank et al., 2008). Moreover, inverting faces generates a larger and sometimes later N170 (Bentin et al., 1996; Eimer, 2000; Eimer et al., 2010; Roxane J Itier and Taylor, 2004; Roxane J. Itier and Taylor, 2004; Rossion et al., 2000; Towler et al., 2012; Watanabe et al., 2003).

Sensory mismatches, like an invalid preview, usually lead to more pronounced neural responses (Dimigen et al., 2012; Näätänen and Kreegipuu, 2011), which is central to current notions of perception (De Lange et al., 2018) and has been interpreted in terms of prediction errors in predictive coding frameworks (Friston, 2010, 2005; Friston and Kiebel, 2009; Garrido et al., 2008; Stefanics et al., 2014). With respect to trans-saccadic perception, the interpretation of the preview effect as a predictive process is particularly intriguing, because a crucial idea for explaining visual stability is that upcoming foveal visual input is predicted based on pre-saccadic peripheral information and a copy of the motor command sent to perceptual brain areas (Cavanaugh et al., 2016; Friston et al., 2012; Melcher and Colby, 2008; Wurtz, 2008).

In Experiment 2, we asked whether the preview effect across a saccade reflects a predictive process across multiple trials. We manipulated the proportion of valid and invalid trials to generate blocks with mostly valid (66.6% valid) and mostly invalid (33.3% valid) trials. Proportion manipulations have successfully demonstrated the predictive nature of sensory processing (Grotheer et al., 2014; Kovács et al., 2012; Mayrhauser et al., 2014; Summerfield et al., 2011, 2008), with the rationale that a more frequent event is more expected than a less frequent event and, therefore, elicits a reduced neural response. Thus, if the preview effect reflected a predictive process, it should become smaller in the mostly invalid and larger in the mostly valid block.

## 2. Materials & Methods

### 2.1. Participants

Twenty volunteers participated in each experiment in return for a monetary reimbursement. All participants gave written informed consent and reported normal or corrected to normal vision that was additionally confirmed by an eye-sight test. In Experiment 1, two participants had to be excluded because of poor performance in the tilt discrimination task. Of the remaining 18 participants, 16 were right-handed, 7 were male, and mean age was 24 years (range: 19-30 years). In Experiment 2, one participant had to be excluded because of bad EEG data resulting from a technical problem during data collection. Of the 19 remaining participants, 16 were right-handed, 6 were male, and mean age was 25 years (range 20-40 years). The procedures of both experiments were approved by the local ethics committee.

### 2.2. Stimuli

Stimuli were presented on a VIEWPixx/EEG monitor (VPixx Technologies Inc., Canada) at 120 Hz screen refresh rate and 1920 × 1080 display resolution. In Experiment 1, 42 face images were taken from the Nottingham face database (http://pics.stir.ac.uk/zips/nottingham.zip) and from the Faces 1999 (Front) dataset (http://www.vision.caltech.edu/archive.html); 21 of which were female and 21 male. In Experiment 2, we selected a set of 16 face images only from the Nottingham face database, of which eight were male and eight female. The face images in this reduced set were more uniform concerning the distribution of facial features across images. For the face images of both experiments, a circular mask with a diameter of 2.88° was centered at the tip of the nose and the image was sized to contain all relevant facial features. The images were placed at 8° eccentricity from screen center. For each original face image, we generated a phase-scrambled counterpart that was presented as a transient (for the duration of 2 frames) during the saccade to achieve the same level of visual change in the display for both valid and invalid preview conditions. The stimuli were processed with the SHINE toolbox (Willenbockel et al., 2010) in Matlab, in order to equate low-level image features that could otherwise present a confound in the EEG signal. We used the function histMatch with the mask option to match the luminance histogram of all face cut-outs and their scrambled counterparts to the average histogram of all face cut-outs within each of the two experiments.

### 2.3. Procedure

Each trial started with a placeholder display consisting of a fixation cross (0.5° by 0.5°) at the screen center and white rings (width 1 pixel) framing the position of the upcoming faces (Figure 1A). In Experiment 1, one white ring appeared on either side of the fixation cross (Figure…), in Experiment 2, only one ring appeared to the left of fixation. Stable fixation within an area of 2° around screen center for 1 second triggered the preview display. In Experiment 1, the preview display contained two faces one at either side from fixation; in Experiment 2, there was only one face to the left of fixation. The face images replaced the placeholder rings. Keeping stable fixation at the center of the preview display for 500 ms triggered the color cue. In Experiment 1, the fixation cross turned either blue or green indicating the saccade direction (color-to-direction assignment counterbalanced across participants). In Experiment 2, the fixation cross turned grey, prompting for a saccade to the single face on the left. Participants had been instructed to respond as quickly and accurately as possible to the cue by making one single eye-movement to the corresponding face stimulus. Saccade onsets were detected online, and upon detection a scrambled version of the preview face was presented for two frames (16.7 ms); in Experiment 1, the faces on both sides were scrambled. The transient occurred no more than 3.5 frames (~30 ms) after saccade onset, with the delay reflecting the computational requirements of saccade detection and the screen refresh rate (Figure 1B). Given a total saccade duration of around 40-60 ms, the target face was presented before fixation onset in most trials (Figure 1C). The purpose of this transient was to roughly equalize the amount of change in the display across all conditions.

**Figure 1.**
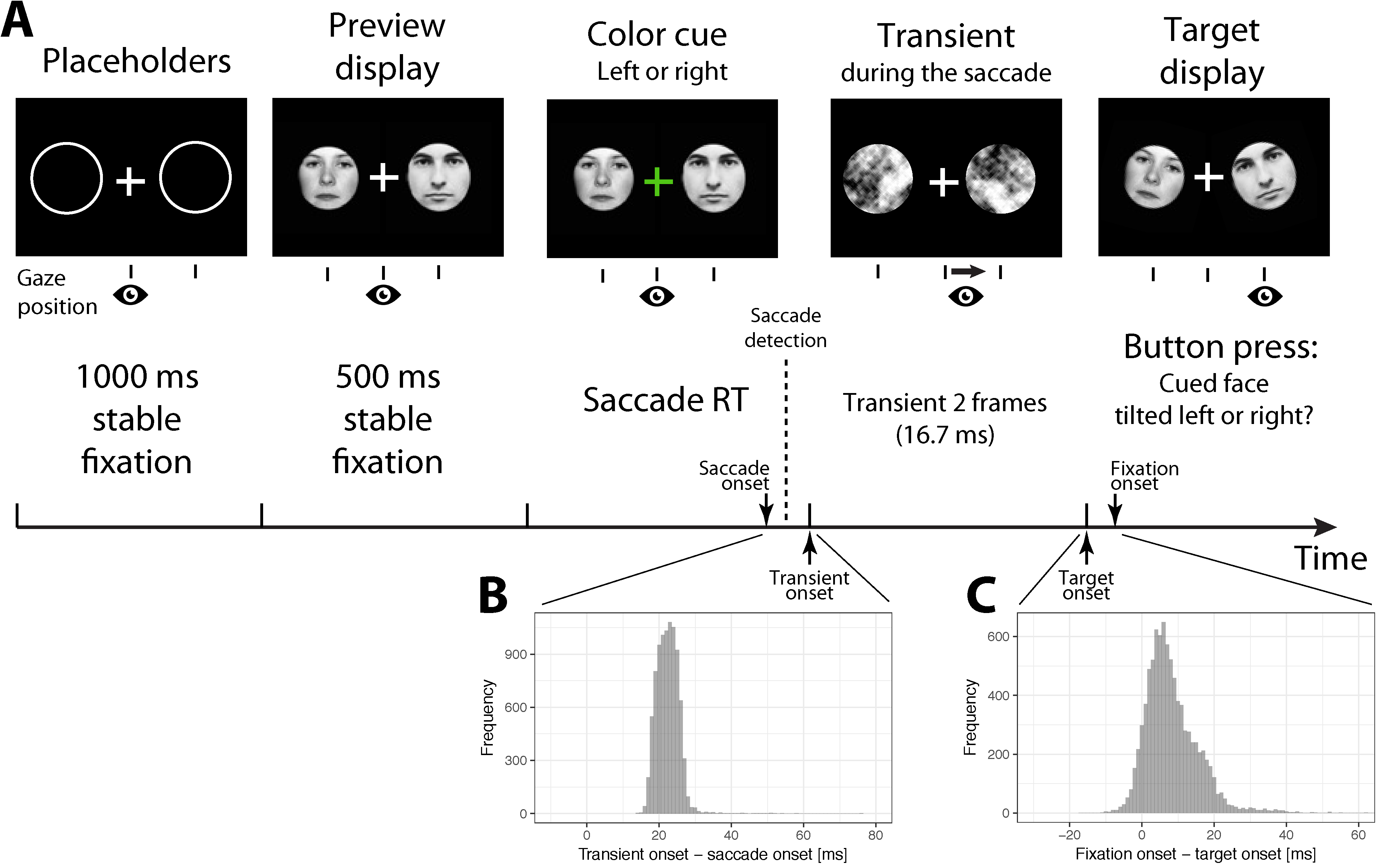
Panel A. Experimental procedure. Stable fixation for 1000 ms triggered the Preview display. Further fixation for 500 ms triggered the color cue indicating saccade direction and, thus, the target face (e.g. green left/blue right, counterbalanced across participants). Both the target (cued) face and non-target face (opposite side) could be either upright or inverted, and could both either change orientation of remain the same across the saccade. After detection of the saccade, scrambled versions of the faces were presented as transients. The speed of saccade detection is illustrated in panel B. The transient was presented most of the time in less than 25-30 ms after actual saccade onset. The transient was replaced by the target display after two frames. The target display contained both target and distractor faces with additional slight tilt (left/right). The target face tilt had to be reported by button press upon fixation onset. The timing of target onset and fixation onset is illustrated in panel C. Fixation onset was most of the time after target onset. Timeline, stimulus size, and target face tilt are not drawn to scale.

During the saccade the faces could change their overall orientation from upright to inverted (or vice versa) or they could remain the same. In Experiment 1, all possible combinations of target and non-target face orientations and changes were realized once with each individual target face, yielding a total set of 672 trials (168 per cell in the crossing of Preview [valid, invalid] and Target Face [upright, inverted] conditions; Figure 2A). In Experiment 2, which employed a smaller set of face images, all possible combinations of target orientations and changes were repeated 16 times for each face. In addition, to investigate whether the preview effect found in Experiment 1 reflected active predictions accumulating across trials, Experiment 2 consisted of two blocks, one containing mostly valid trials (66.6% valid, 33.3% invalid) and the other one containing mostly invalid trials (33.3% valid, 66.6% invalid) (Figure 2B). We expected the preview effect - the difference in the dependent variable between invalid minus valid trials - to be larger in the mostly valid block and smaller in the mostly invalid block (Figure 3). The order of blocks was counterbalanced across participants.

**Figure 2.**
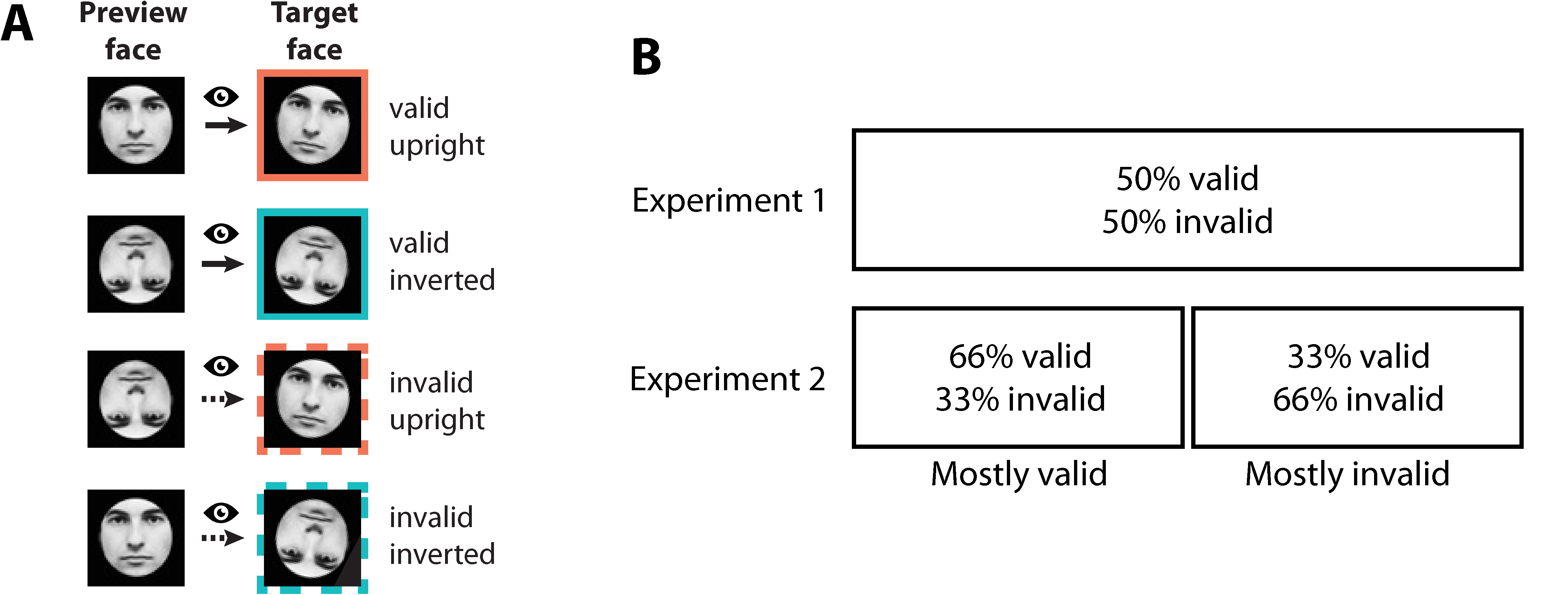
Panel A shows the four possible preview and target face orientation conditions. Both preview orientation and target orientation could be upright or inverted leading to in total four conditions, two of which contained a valid preview (preview orientation and target orientation matched) and two an invalid one (preview orientation and target orientation did not match). Panel B shows the proportion of valid and invalid trials in Experiment 1 and 2. In Experiment 1, valid and invalid trials occurred at a frequency of 50% throughout the experiment. Experiment 2 consisted of two blocks, one with mostly valid (66.6% valid, 33.3% invalid) and one with mostly invalid trials (33.3% valid, 66.6% invalid). Block order was counterbalanced across participants.

**Figure 3.**
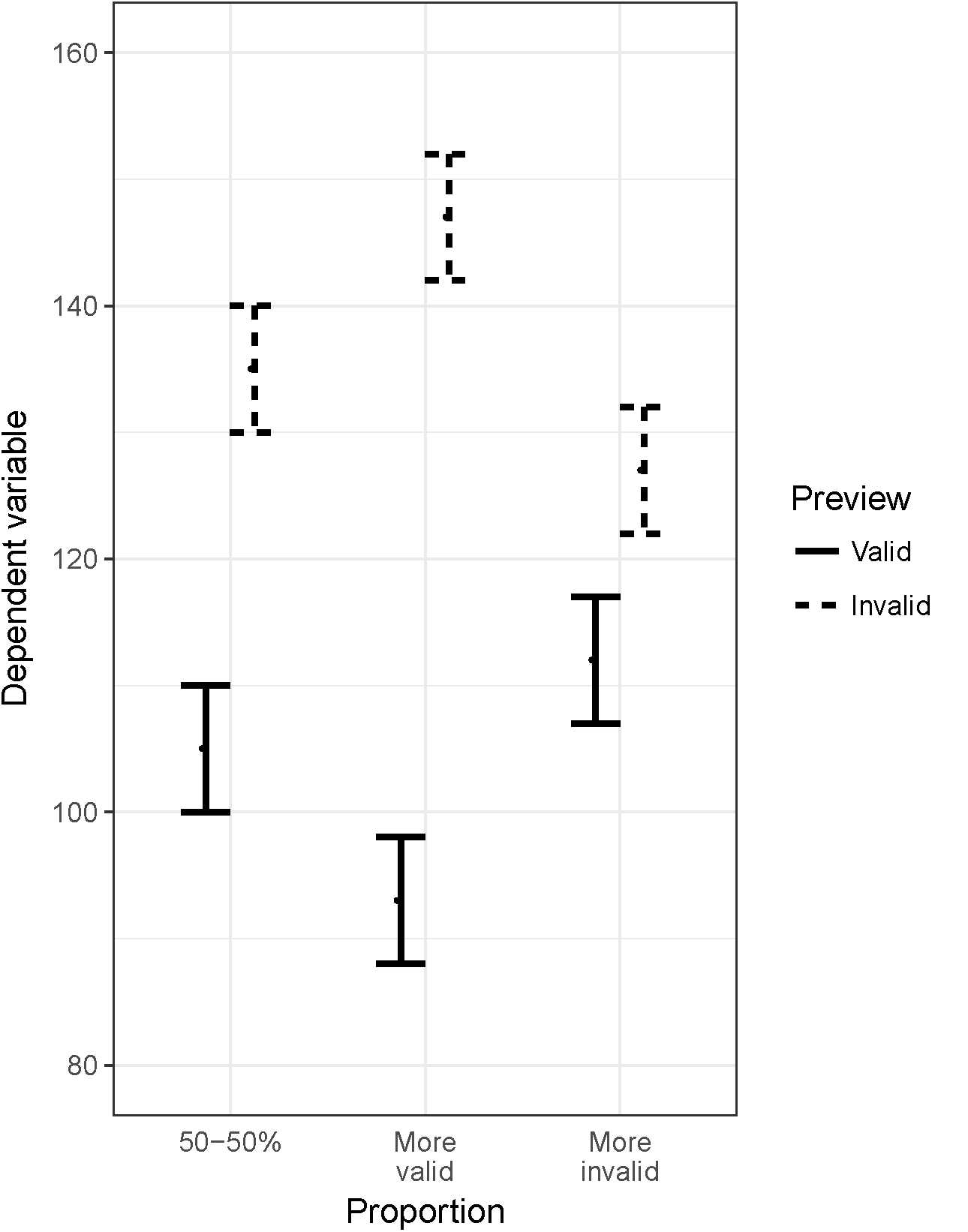
Illustration of the logic of the proportion manipulation to determine the predictive nature of the preview effect (difference on the y-axis between valid, solid, and invalid, dashed, conditions). If the preview effect is predictive, a block with more valid trials is expected to increase the preview effect, and a block with more invalid trials is expected to decrease the preview effect.

Experiment 2 was thus composed of 1024 (with either 171 or 85 per cell in the crossing of Preview [valid, invalid], Target face [upright, inverted], and Proportion [mostly valid, mostly invalid] conditions). For instance, in the mostly valid block, there were 171 valid trials with target upright, 171 valid trials with target inverted, 85 invalid trials with target upright, and 85 invalid trials with target inverted. Importantly, the proportion manipulation was not mentioned to the participants at any point.

In addition to its main orientation (upright or inverted), each target face was slightly tilted (1.8°) either to the left or right, counterbalanced across trials. The non-target face in Experiment 1 had the same amount of tilt as the target face (on the other side of fixation), but its direction (left or right) was random. The target face tilt direction had to be reported by the participants via a computer keyboard with the left and right index finger after they had made an eye-movement to the target face. The purpose of the tilt discrimination task was to ensure that participants paid attention to the target face and gave a response that was orthogonal to all experimental manipulations. In fact, the preview images were not tilted, making them task-irrelevant for the perceptual tilt discrimination response. Correct saccades (end point at least within 2.16° of target face center) were detected online, and participants received feedback in case of incorrect response or if the recorded gaze position was too far from the expected saccade start or end locations. The eye-tracker was recalibrated when it failed to correctly track gaze position, meaning that the experiment did not advance automatically although the participant was adhering to the instructed gaze procedure.

### 2.4. EEG and eye-tracking data recording

The electroencephalogram (EEG) was recorded from 62 electrodes placed at a subset of the locations of the 10-10 system. The right mastoid served as online reference.

Eye-movement data was recorded by an Eyelink 1000 video-based eye-tracker (SR Research, Ontario, Canada) in the desktop mount setup. Default settings for saccade detection were used (velocity threshold 35°sec, acceleration threshold 9500°sec2). The online saccade detection that triggered the scrambled transient (see Procedure) was, however, based on a custom-made algorithm, since the default saccade start events were not transferred quickly enough from the eye-tracking host computer to the experiment workspace in Matlab. We set the heuristic_filter option of the eye-tracker to level 2 in order to receive cleaner gaze position data, despite the minimal additional delay introduced by the higher filter level. A gaze position difference of 0.18° between two subsequent samples, converted to screen pixels depending on individually measured viewing distance of each participant, triggered presentation of the scrambled transient at the next possible screen refresh. This procedure resulted in quick and satisfactory saccade detection in most of the trials (cf. Figure 1B). Both eye-tracking and EEG data were recorded at 1000 Hz sampling rate. Trigger signals were sent to both data acquisition systems by means of a parallel port splitter cable. The trigger signals were used offline to synchronize both data streams for subsequent analysis.

### 2.5. EEG and eye-tracking data analysis

EEG and eye-tracking data was processed in Matlab with the EEGLAB (Delorme and Makeig, 2004) and CoSMoMVPA (Oosterhof et al., 2016) toolboxes. The eye-tracking data was synchronized with the EEG by means of the eyeeeg plugin (Dimigen et al., 2011). Upon synchronization, the signal was down-sampled to 250 Hz, low-pass filtered (Hamming windowed sinc FIR filter, 40 Hz, transition band width 10 Hz, cutoff frequency [-6 dB] 45 Hz), and re-referenced to average reference (Hinojosa et al., 2015). The EEG data was then visually inspected for major artifacts. Portions of data with severe artifacts were removed and bad channels were spherical-spline interpolated.

In order to correct eye movement artifacts in the EEG, we applied independent component analysis (ICA; Makeig, Bell, Jung, & Sejnowski, 1996). Eye-movement related components were determined based on the variance ratio of component activation during periods of eye-movements (blinks and saccades) versus periods of fixations (Plöchl et al., 2012). ICA was conducted in a separate processing pipeline containing an additional high-pass filter (Hamming windowed sinc FIR, 1 Hz, cutoff frequency [-6 dB] 0.5 Hz) that was applied after down-sampling and before low-pass filtering. The ICA algorithm was infomax (Bell and Sejnowski, 1995) with the pca option to ensure proper rank of the data matrix. The ICA results (sphere and weights) were transferred to the corresponding datasets in the original processing pipeline without the severe high-pass filter. Components were then rejected if the mean variance of their activation across eye-movement periods was 10% greater than the mean variance across fixation periods.

In both experiments, we extracted epochs of interest for periods of target fixation. Target fixation epochs were extracted from −200 to 600 ms with respect to target face fixation onset. Baseline correction was conducted with respect to the 200 ms period before onset of the preview display. This approach was adopted for two reasons: first, to compare the post-saccadic activity to a period in which there was no visual input, and, second, to prevent possible residual eye-movement-related activity from confounding the baseline correction. In Experiment 2, we also extracted epochs of interest for the time of preview display, from - 200 to 800 ms with respect to preview display onset, with baseline correction for the 200 ms prior to preview display onset.

Only trials with correct responses and trials in which participants had followed the gaze instructions in the experimental procedure were included in the analysis. These were trials in which participants kept stable fixation within 2° of screen center, made no saccades before cue onset, and the saccade end point had to be within 2.16° of target face center. If the target had not been presented before fixation onset, due to a delay in saccade detection, the time difference between fixation onset and target onset had to be less than 20 ms (see Figure 1C and Procedure for details), which is largely within the time course of saccadic suppression (Benedetto and Morrone, 2017; Bremmer et al., 2009; Diamond et al., 2000). This restriction was disregarded in Experiment 2 for the preview-locked analysis only, because this analysis focused on the time period before the saccade and disregarding this criterion increased the number of available trials. Finally, trials with very fast and very slow responses in the tilt discrimination task were excluded by a median absolute deviation filter with a conservative criterion of 3 (Leys et al., 2013). In Experiment 1, these strict criteria led to acceptance of a median number of 104 trials ranging from 58 to 139 across participants and cells of the design preview by target orientation. In the fixation-locked analysis of Experiment 2, median number of accepted trials was 78, ranging from 32 to 165, across cells of the same design extended by the factor proportion. For the preview-locked analysis of Experiment 2, the median number was 79, and the range was the same. The extended range in Experiment 2 compared to Experiment 1 was due to the proportion manipulation which lead to an unbalanced number of trials across cells of the design.

To determine whether and how the pre-saccadic preview affected processing of the post-saccadic target face, we investigated the time course of Preview orientation (upright, inverted) and Target orientation (upright, inverted) effects in the EEG with a whole-scalp Bayes factor analysis. ERP components are known to differ across tasks, and since we used a novel gaze contingent task, such an analysis reduces the risk of false positive findings (Luck and Gaspelin, 2017). Note, that the same conditions resulting from the factors Preview orientation (upright, inverted) and Target orientation (upright, inverted) can be modelled equally well by either of the factors Target or Preview orientation (upright, inverted) together with a Preview factor (valid, invalid) which indicates whether the target and the preview face were of the same (valid) or different (invalid) orientation.

Experiment 1 included the factor Cue direction (left, right; synonymous with saccade direction) and, for lateral electrodes, also the factor Laterality (contra, ipsi; with respect to cue direction). To create the Laterality factor, EEG data from trials with saccades to the left were swapped across hemispheres in order to assign left hemisphere electrodes to the contralateral, and right hemisphere electrodes to the ipsilateral condition. For instance, the signal at electrode PO7 was assigned the label *ipsilateral* for leftward saccade trials and the label *contralateral* for rightward saccades trials. The signal at electrode PO8 was treated in the opposite way. In contrast to Experiment 1, Experiment 2 omitted the factors Cue direction and Laterality, because there was only one target face to the left to which saccades were directed, but instead it included the factor Proportion (mostly valid, mostly invalid). For Experiment 2, we additionally analyzed the data time-locked to the preview display in order to determine any pre-saccadic expectation effects introduced by the proportion manipulation.

The preview-display locked analysis of the EEG data revealed an interesting unexpected result, with the face inversion effect in the N170 triggered by the preview display occurring later than the face inversion effect triggered by the target display. We tested the reliability of this delay by analyzing onset latencies of the N170 face inversion effect. Since this was a post-hoc analysis, this result might be less reliable.

In addition to the whole-scalp Bayes factor, we also computed common repeated measures Anovas on average ERPs at selected electrode sites and for time-windows of main interest to further consolidate the results.

### 2.6. Whole-scalp analysis

At each electrode and time point, we computed a Bayes factor (BF) based on the average EEG voltage across trials per participant and condition. We used the BayesFactor package (version 0.9.12-2) in R (R Core Team, 2013) with fixed-effect priors set to the default Cauchy distribution at location 0 and scale 0.5. This prior can be verbally expressed as expectation of a medium-sized effect with smaller effects being more likely than larger effects (Rouder et al., 2009). In contrast to null-hypothesis significance testing, the Bayes factor provides a measure of graded evidence for the presence versus absence of an effect (Dienes, 2016; Rouder et al., 2016; Wagenmakers, 2007). In line with common practice, we consider a BF greater than 3 as positive evidence, a BF lower than 1/3 as negative evidence, and a BF between 1/3 and 3 as non-decisive (Raftery, 1995).

To obtain a BF for a main or an interaction effect in a multifactor design, such as in the present study, it is advisable to calculate the so-called BF *across matched models.* This is because the BF is a likelihood ratio that results from comparing two models, which is usually the likelihood of the data given the alternative hypothesis/model divided by the likelihood of the data given the null hypothesis/model. A multifactor design offers many pairs of models with one model containing the effect of interest and the other not. Thus, there are many possible likelihood ratios which could be considered as providing the BF for a certain effect.

The most straightforward way to solve this problem is to compute the sum of the likelihoods of all of the models with the effect of interest and divide it by the sum of the likelihoods of all of the corresponding models without the effect of interest. Models containing higher-order interactions with the effect of interest are disregarded. This procedure is, for instance, implemented in the software JASP (JASP Team, 2018).

## 3. Results

### 3.1. Experiment 1: Valid peripheral preview improves post-saccadic tilt discrimination performance

We analyzed manual response times in the tilt discrimination task only for those trials that entered the EEG analysis, which also excludes tilt discrimination errors. Error trials were, however, included in the error rate analysis, which still excluded trials with incorrect saccades (see Methods). For both computations the design contained three factors: Target Orientation (upright, inverted), Preview (valid, invalid), and Cue Direction (left, right; equivalent with saccade direction).

As expected, a valid preview led to on average shorter response times than an invalid preview (valid 1,200 ms, invalid 1,229 ms), *F*(1,17) = 16.26, *p* = .001, BF = 13.20 (Figure 4A) which is in line with the behavioral preview benefit effect in reading research (Rayner, 1975; for a review see Schotter et al., 2012). Error rates were the same in both preview conditions (valid 17 %, invalid 18 %), *F*(1,17) = 0.80, *p* = .382, BF = 0.24 (Figure 4B). Performance was also affected by target face orientation. Upright target faces led to a faster response than inverted target faces (1,186 ms versus 1,243 ms), *F*(1,17) = 24.31, *p* < .001, BF > 100. Upright faces were also less error prone (15 %) than inverted ones (20 %), *F*(1,17) = 21.97, *p* < .001, BF > 100. This effect was, however, not of primary interest in the current study.

**Figure 4.**
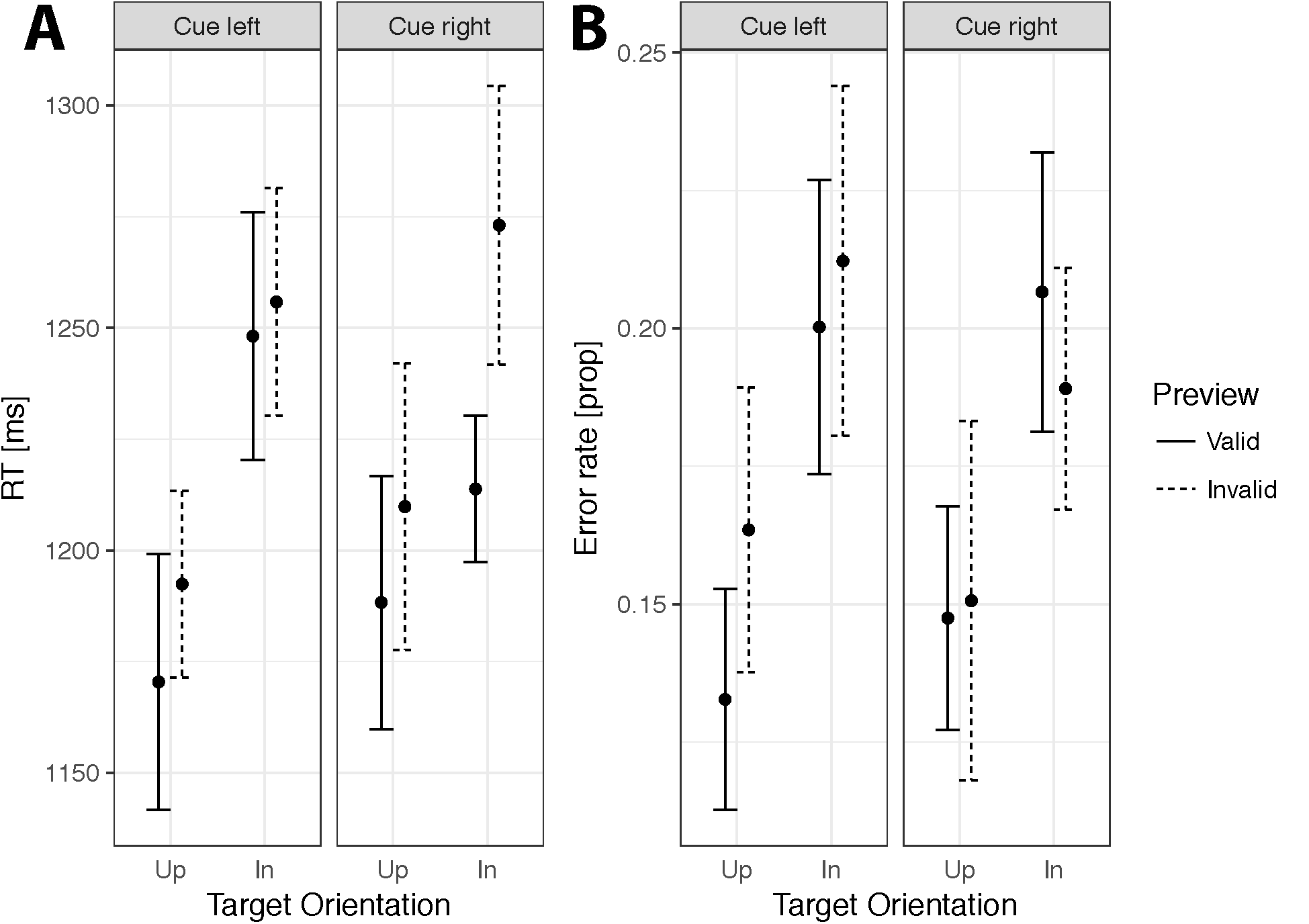
Mean response times (panel A) and error rates (panel B) in the tilt discrimination task in Experiment 1, split by the factors Cue Direction, Target Orientation, and Preview. Participants were faster in valid (solid) than in invalid preview conditions. Target orientation also affected the response: Participants responded faster (panel A) and made fewer errors (panel B) in trials with upright (Up) compare to with inverted (In) target faces.

The ANOVA also suggested that the Preview effect might have been influenced by Cue Direction, with a marginally significant interaction in response times, *F*(1,17) = 4.29, *p* = .054, and significant interaction in error rates, *F*(1,17) = 10.56, *p* = .005. In response times, this effect indicated a larger preview effect for right side targets; for error rates it indicated the opposite pattern. However, the BF for response times was BF = 0.65 and for error rates it was BF = 0.79, suggesting that strong conclusions should not be drawn from these results.

### 3.2. Experiment 1: Valid peripheral preview reduces the fixation-locked N170 (fN170) amplitude

The results of the fixation-locked whole-scalp Bayes factor analysis are illustrated in Figures 4 and 5. Figure 5 shows the BF for the theoretically most relevant effects of Preview Orientation (panel A, aka Preview x Target Orientation interaction), Target Orientation (panel B), and the Preview effect (panel C, aka Preview Orientation x Target Orientation interaction). The ERPs corresponding to these effects are illustrated in panel D. Note that the Preview Orientation (upright, inverted) main effect is expressed as a Preview x Target Orientation interaction.^1^

**Figure 5.**
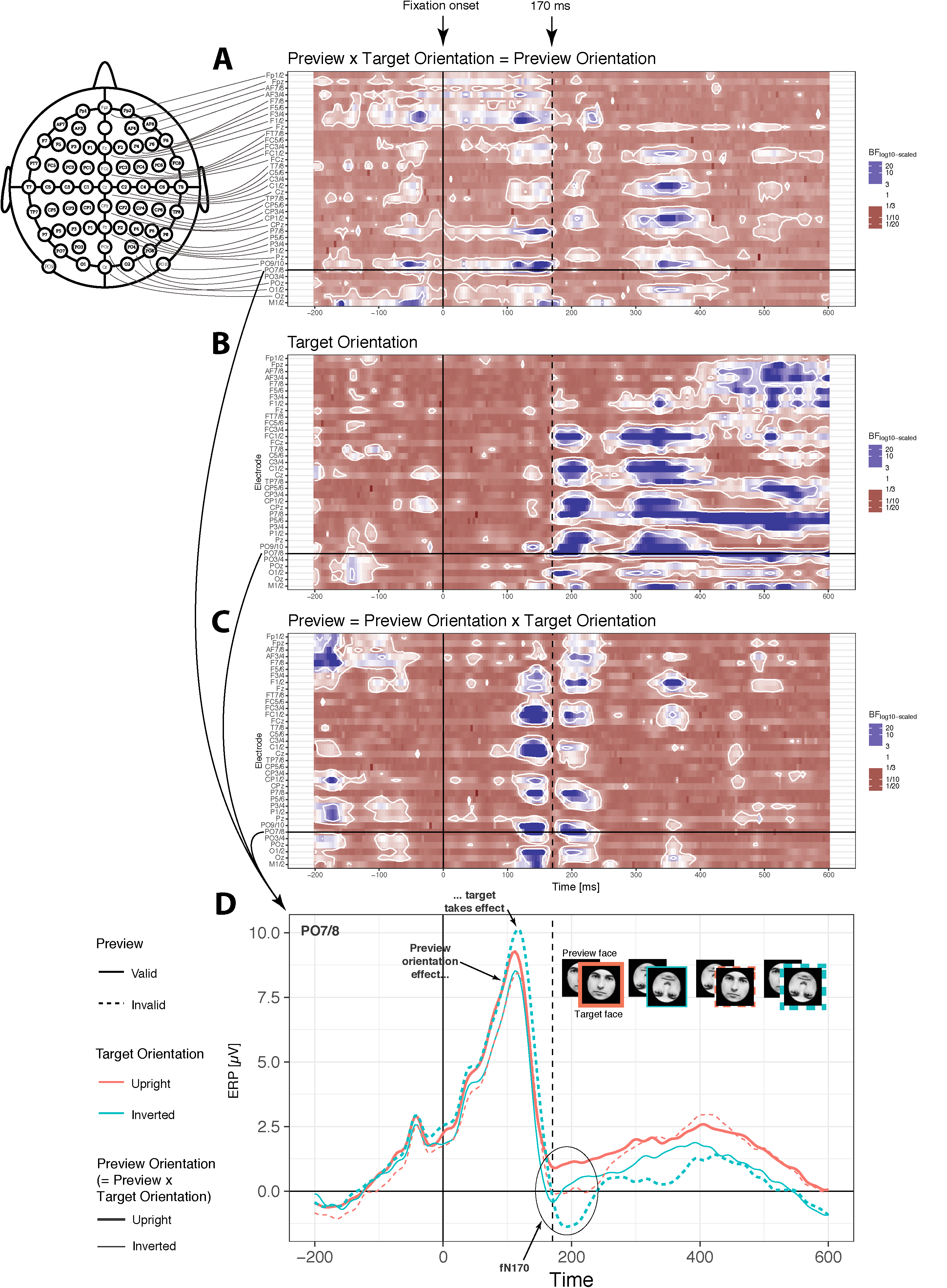
Whole-scalp Bayes factor (BF) analysis of EEG data time-locked to fixation-onset on the target face (panels A-C). Panel D illustrates the corresponding ERPs at electrode pair PO7/8. Each horizontal row of panel A-C represents the time-course of the BF for one contra-ipsilateral electrode pair, sorted from frontal (top) to posterior (bottom) sites and within this order further from lateral (top) to medial (bottom) sites. Values greater than 3 (blue) denote positive evidence, values less than 1/3 (red) negative evidence. Values in-between are indecisive (white). The thresholds 3 and 1/3 are indicated by two-dimensional white contour lines. The vertical dashed line at 170 ms only serves as visual guide and does not indicate any event in the experiment. Panel A shows the Preview x Target Orientation interaction, aka Preview Orientation main effect. From ca. 100 ms post fixation onset to 170 ms the orientation of the preview face dominated the posterior lateral EEG signal (see also panel D). Evidence for this effect became positive again between ca. 300 to 400 ms primarily at central-parietal sites. Panel B illustrates the main effect of Target Orientation. Evidence for this effect became positive from ca. 170 ms post fixation-onset at lateral posterior and some central sites and, after some decrease in evidence from ca. 250 to 300 ms extended throughout the post-saccadic time-window. The corresponding face inversion effect in the fN170 is illustrated in panel D. Panel C shows evidence for the crucial Preview effect, aka Preview Orientation x Target Orientation interaction. In time windows of ca. 50 ms before and after 170 ms the EEG response was more pronounced in valid (preview orientation and target orientation matched) compared to invalid (no match) conditions. The ERPs in panel D show this effect in the fN170 component at electrode pair PO7/8. Note that baseline correction was conducted with respect to the time window −200 to 0 ms before preview display onset which is outside the plotted time period (cf. Figure 1).

Interestingly, as can be seen from Figure 5, the initial phase of the fixation-locked EEG response already showed some evidence for an influence of the orientation of the preview face (panel A), which became decisively positive (BF > 3, color-coded in blue within white contour lines) from around 110 to 170 ms post fixation onset. During this relatively early period after fixation onset the preview face was no longer presented on the screen but instead had been replaced by the target face, which could have had a different orientation than the preview face. Nevertheless, an inverted preview face led to a more negative EEG response than an upright preview face (see panel D), which perhaps indicates that a late face processing sensitive component, such as the N250 or N400, carried over from the pre-saccadic period. This effect is quite interesting because it could reflect a mechanism relevant for the experience of visual stability. Immediately after the fixation, the EEG signal initially reflects what we perceived before the saccade and expect to see after the saccade, until new post-saccadic information is incorporated (Mirpour and Bisley, 2016). For face orientation this updating process apparently happens at around 170 ms. Indeed, the switch at 170 ms is consistent with the timing of the face-selective N170 component.

Almost exactly at 170 ms the main influence on the EEG signal switched from the preview face to the target face (cf. Figures 5A and 5B), which elicited a more negative response for inverted than for upright target faces (Figure 5D). This modulation perfectly matches the classic N170 face inversion effect (Bentin et al., 1996; Eimer, 2000; Eimer et al., 2010; Roxane J Itier and Taylor, 2004; Roxane J. Itier and Taylor, 2004; Rossion et al., 2000; Towler et al., 2012; Watanabe et al., 2003). We consider this target orientation effect around 170-220 ms post fixation as a modulation of the fixation-locked N170 component, the fN170. Most importantly, for a period of about 80 ms before and after the crucial time point of 170 ms, the preview orientation and target orientation factors interacted (Figure 5C), showing a more pronounced neural response when the preview face and target face orientations matched (valid preview) compared to when they did not match (invalid preview) (Figure 5D). As can be seen from Figure 5D, the fN170 component in particular was more pronounced in invalid (dashed lines) than in valid preview (solid lines) conditions, which is consistent with the idea of a trans-saccadic prediction error. The role of prediction was further explored in Experiment 2.

As can be seen from Figure 5D, Preview effect and Target Orientation interacted again from around 320 ms post fixation for a duration of about 80 ms in particular at central parietal electrodes. The target orientation effect here consisted in a more negative deflection for inverted compared to upright target faces and this face inversion effect was larger for invalid than for valid preview conditions. This interaction probably reflected increased processing of the target face orientation in invalid than in valid preview conditions, which appears intuitively plausible. With an invalid preview, the target face presented new information which requiring more in-depth processing of the critical feature face orientation.

As can be seen from Figure 6, Preview and Target Orientation factors did, with one exception (three-way interaction with Cue Direction, Figure 6H), not interact with other factors. This interaction with Cue Direction showed sufficient positive evidence before and around the time of the saccade and suggested that the Preview x Target Orientation interaction, aka Preview Orientation main effect consisted in a more negative EEG for inverted compared to upright preview faces, which was more pronounced for cue/saccade right trials than for cue/saccade left trials (direction of effects not illustrated). Given the posterior lateral distribution of this effect (electrodes O1/2, PO9/10), and the time periods before and around the time of the saccade, this effect might have reflected saccade-related perceptual processes.

**Figure 6.**
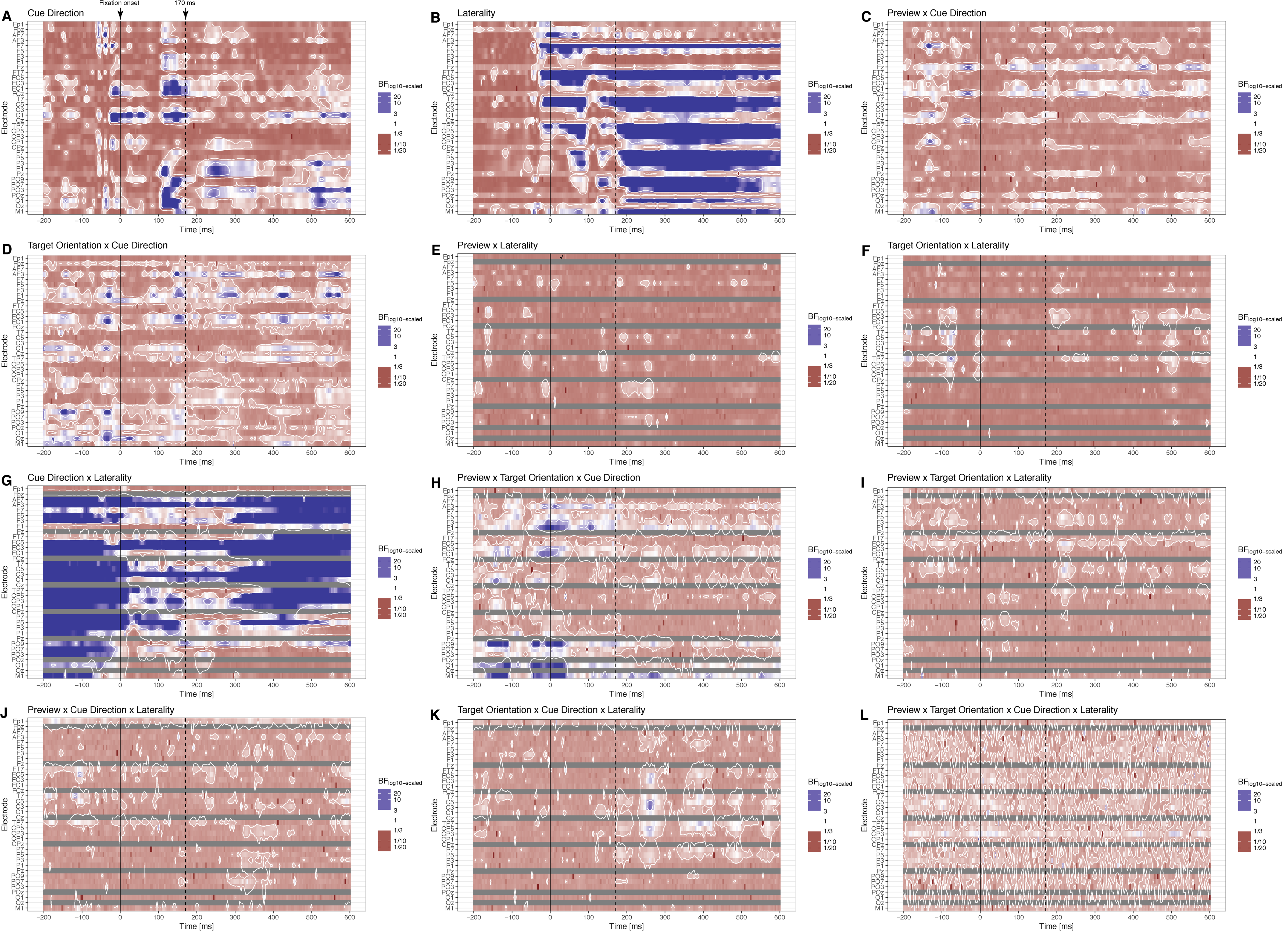
Whole-scalp Bayes factor (BF) for all the remaining main and interaction effects of Experiment 1 not illustrated in Figure 5. Importantly, the Preview and Target Orientation effects did not interact with other factors in particular not in the spatio-temporal window of the fl70 preview effect at lateral posterior electrodes ca. 50 ms before and after the 170 ms time stamp.

Additional effects of less theoretical significance were identified in our analyses, including a main effect of Cue Direction (Figure 6A), and the substantial effects of Laterality (Figure 6B) as well as the Laterality x Cue Direction interaction (Figure 6G). The Cue Direction effect indicated evidence for differences between right side and left side saccade trials at posterior lateral electrodes from ca. 100 to 160 ms and at central electrodes from during the saccade to 170 ms post fixation (Figure 6A). The Laterality effect showed strong evidence for widespread effects across the whole post-saccadic time period (Figure 6B). Finally, Laterality and Cue Direction showed a pronounced interaction across several electrode sites and across the whole analysis time window (Figure 6G). Such laterality effects might be related to face processing differences between hemispheres (Frässle et al., 2016; Schweinberger et al., 2004) or some of fact which is not particularly central to the current study. These factors were modeled in the analysis in order to control for potential interactions with the preview and target orientation effects, which were of more central theoretical interest.

### 3.3. Experiment 1: Anova on average ERPs in the fN170 time window in line with the whole-scalp analysis

To provide a statistical assessment of the main results from a frequentist perspective, we computed repeated measures Anovas on average ERPs at electrode pair PO7/8, which is known to show the most pronounced N170 effects (Hinojosa et al., 2015), in the time window from 165 to 250 ms. This time window is later than the usual time window in ERPs studies on the N170 effect (Bentin et al., 1996), but seems to be more appropriate given the extended N170 in the invalid preview conditions in our experiment (cf. Figure 5). To assess the later central-parietal Preview x Target Orientation interaction, we additionally computed a repeated measures Anova at electrode CPz for the later time window of 320 to 400 ms. The Anova results were in line with the evidence from the whole-scalp BF analysis. The Anova showed clear main effects of Preview, *F*(1,17) = 36.55, *p* < .001, and Target Orientation, *F*(1,17) = 8.50, *p* = .010, which corroborated the more pronounced N170 in invalid compared to valid preview conditions and the more pronounced N170 for inverted compared to upright target faces. The Target Orientation x Cue Direction interaction was almost significant, *F*(1,17) = 4.01, *p* = .062, but the corresponding BF = 0.30 suggested that the evidence for this effect is negative. We will, therefore, not consider this effect any further. There was also a clear effect of Laterality, *F*(1,17) = 20.16, *p* < .001, indicating a more negative ERP contralateral to the side of the target face.

One effect markedly differed between the Anova on average ERPs and the whole-scalp BF analysis. The Anova showed a highly significant Preview x Laterality interaction, *F*(1,17) = 21.53, *p* < .001, with, however, a very low BF = 0.33 calculated on the same values (see also Figure 6E) indicating that this interaction did not have an effect. This discrepancy between frequentist and Bayesian results suggests that the effect is not very reliable, although it would have been theoretically meaningful. The direction of the interaction suggested a larger preview effect, i.e. difference between valid and invalid trials, at electrodes contralateral to target/saccade side compared to ipsilateral electrodes. If anything, one would have expected this direction of the effect, because the contralateral hemisphere is the hemisphere to which the preview stimulus is projected.

The Anova at electrode CPz on average amplitudes for the 320 to 400 ms time window confirmed the Preview x Target Orientation interaction, *F*(1,17) = 10.68, *p* = .005, and corroborated the more pronounced target face inversion effect (upright minus inverted) with an invalid (−1.19 μV) compared to with a valid (−0.07 μV) preview. This Anova also showed a main effect of Target Orientation, *F*(1,18) = 5.90, *p* = .027. No other effects were statistically significant.

### 3.4. Experiment 2 replicates the effects from Experiment 1 in tilt discrimination performance and in the fixation-locked EEG

In contrast to Experiment 1, Experiment 2 contained a more restrictive selection of face stimuli, which were only presented to the left of fixation and the proportion of valid and invalid trials was manipulated to achieve a mostly valid (66.6% valid, 33.3% invalid) and a mostly invalid (33.3% valid, 66.6% invalid) block. Overall, Experiment 2 replicated the preview effects in both behavioral (Figure 7) and fixation-locked EEG data (Figure 8). Response times in the tilt discrimination task were faster in valid than in invalid preview conditions, *F*(1,18) = 31.58, *p* < .001, BF = 4.89 (Figure 7A). There was no preview effect in error rates *F*(1,18) < 1, BF = 0.19 (Figure 7B). The fixation-locked EEG exhibited again a pronounced preview effect in the fN170 component (Figure 8E), which was corroborated by a repeated measures Anova on average ERPs at right hemisphere electrode PO8 in the time window 165 to 250 ms, *F*(1,22) = 41.46, *p* < .001. Note that, since preview face stimuli were only presented in the left visual field in this experiment, we focused the ERP analysis on the right hemisphere, that is at posterior-lateral electrode PO8. The evidence for the preview effect was, however, similar at the corresponding electrodes on the left hemisphere as can be seen from Figure 8E.

**Figure 7.**
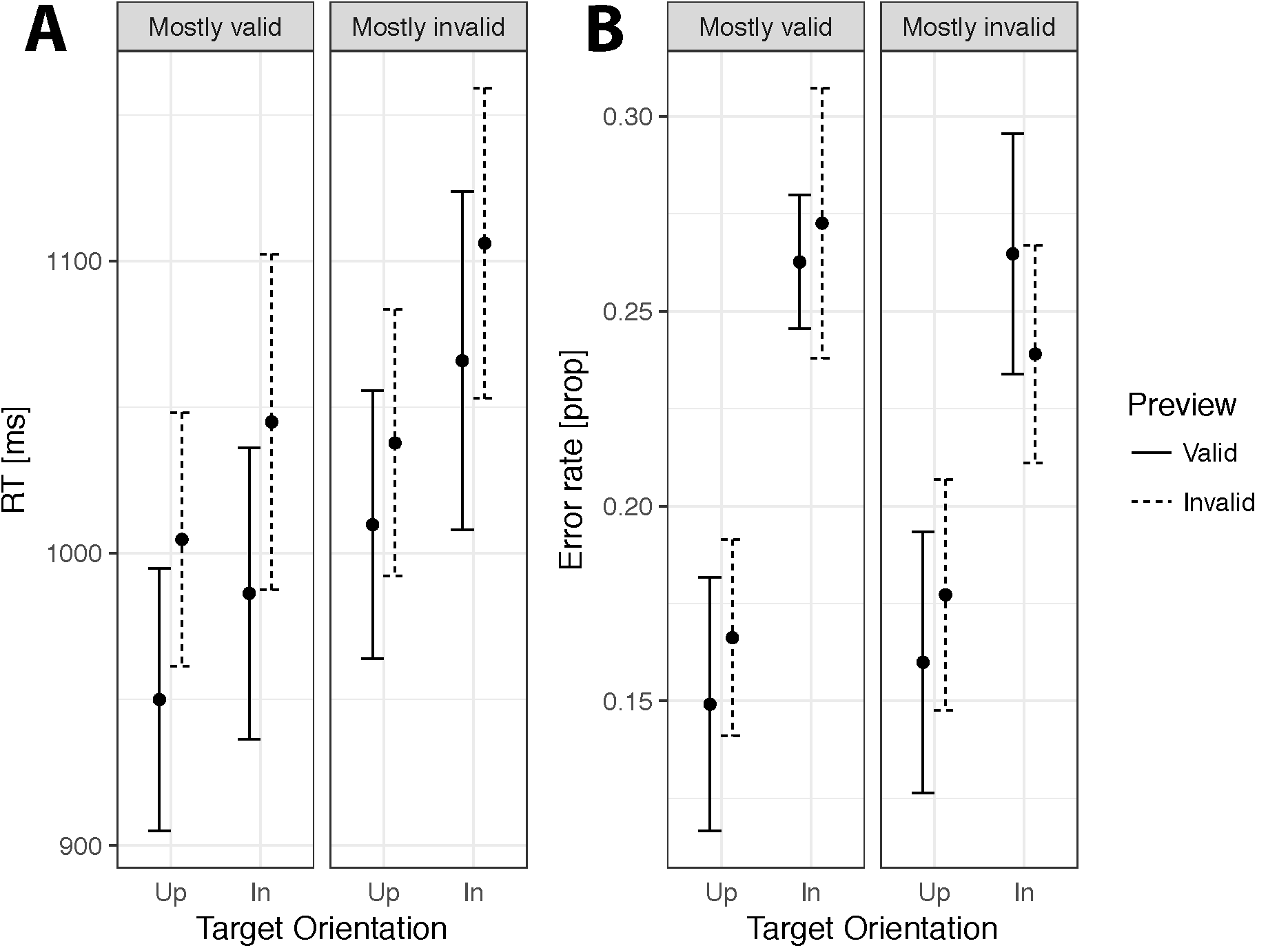
Behavioral results of Experiment 2. Response times (panel A) were faster in valid than in invalid trials, and faster for upright (Up) than for inverted (In) targets. The evidence for the Preview (valid, invalid) by Proportion (mostly valid, mostly invalid) interaction was unclear (see text). Error rate (panel B) was lower for upright than for inverted targets.

**Figure 8.**
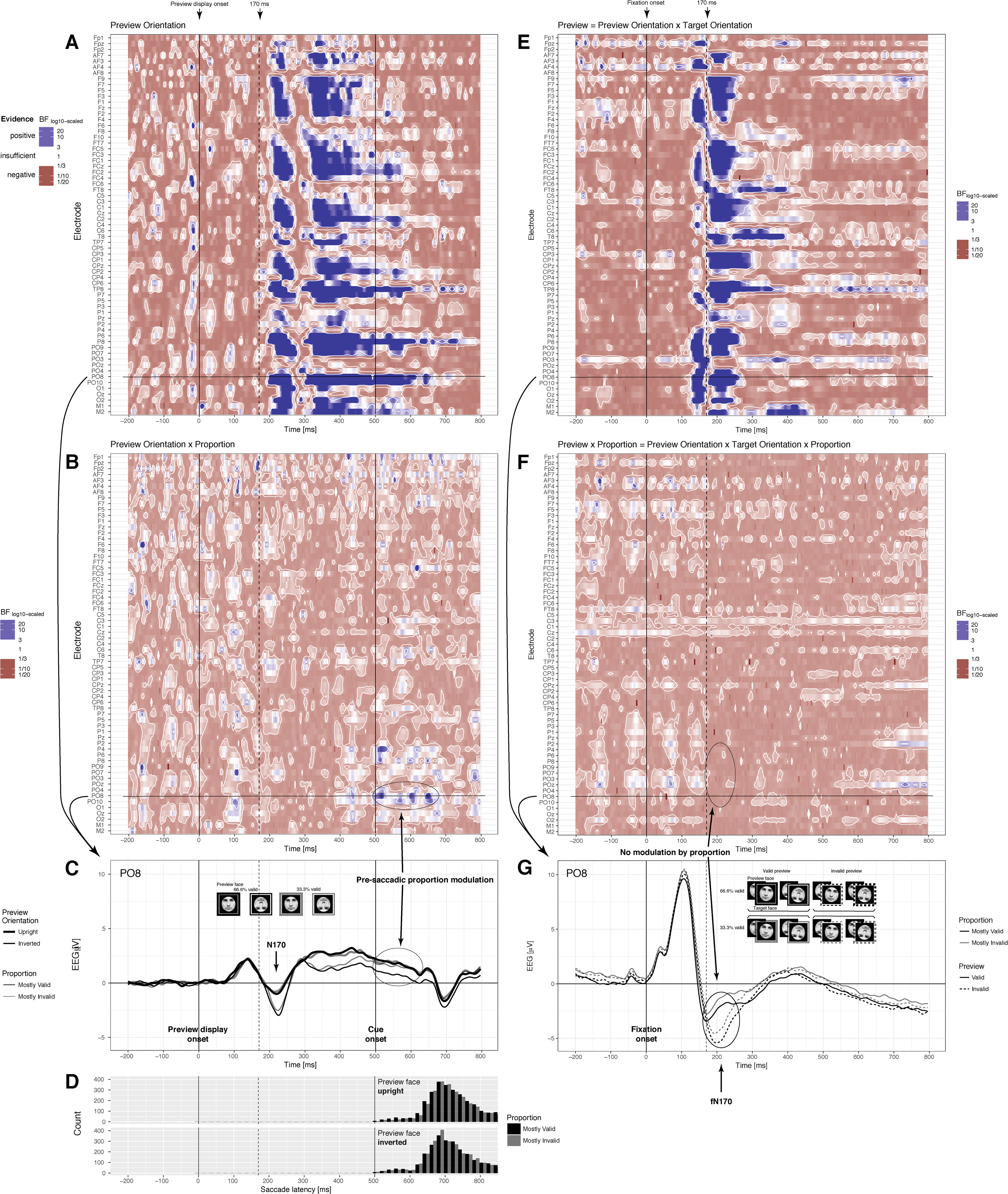
Whole-scalp Bayes factor (BF) for Experiment 2, time-locked to preview display onset (panels A-C), histograms of saccade latencies (panel D), and time-locked to fixation-onset on the target face (panels E-G). The preview period (panel A) showed positive evidence for a Preview Orientation effect in the N170 and in a later component from ca. 300 ms in with more negative deflections for inverted faces (panel C). With cue onset and before onset of most of the saccades (pane D) this face inversion effect at posterior lateral electrodes disappeared earlier in the mostly invalid than in the mostly valid block (panel C) as evidenced by a Preview Orientation x Proportion interaction (panel B). The preview effect in the fN170 established in Experiment 1 was replicated in Experiment 2 (panel E). Crucially, the fN170 preview effect was the same in mostly valid and mostly invalid blocks (panel G) as evidenced by a BF clearly lower than 1/3 for the Preview x Proportion interaction (panel F). Panel G contains ERPs averaged across both target orientations (upright, inverted). For effects of target orientation see Figure 9. Note that baseline correction was conducted for the −200 to 0 ms time window before preview display onset (panel C).

**Figure 9.**
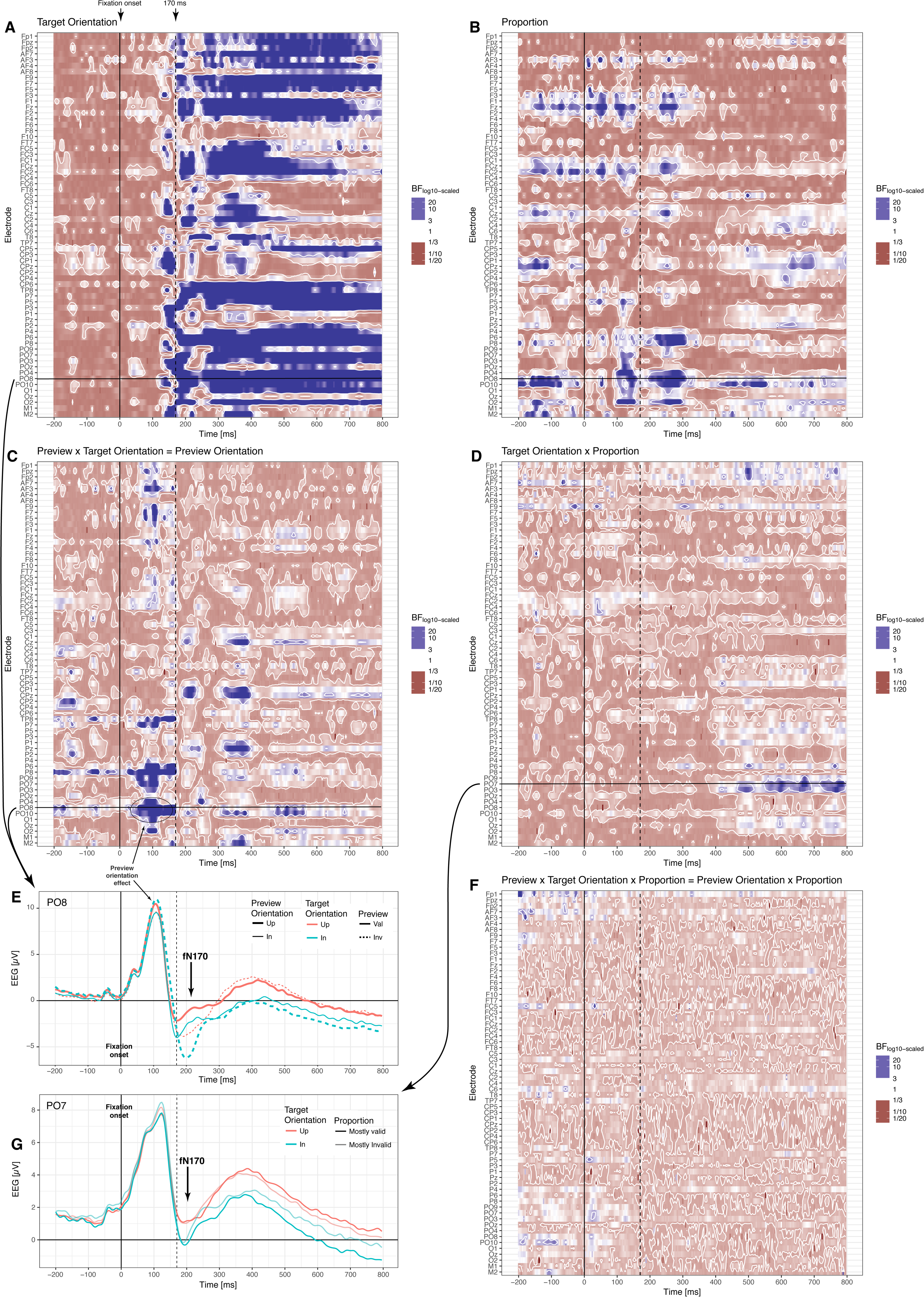
Fixation-locked whole-scalp Bayes factor (BF) for the remaining main and interaction effects of Experiment 2 not illustrated in Figure 8. The effects of Experiment 1 were replicated. Target Orientation elicited again a pronounced face inversion effect in the fN170 and a later component commencing at ca. 300 ms post-fixation onset (panel A, panel E). Preview Orientation showed again a face inversion effect in the initial phase of post-saccadic processing before 170 ms after fixation onset (panel C, panel E). In addition, the evidence for a more negative fN170 in mostly valid compared to mostly invalid blocks was clearly positive (Proportion main effect, panel B, corresponding ERPs in Figure 8G). The Target Orientation effect was more sustained in the mostly valid compared to the mostly invalid blocks in a very late time window and surprisingly at ipsilateral sites (panel D). Evidence for the three-way interaction was largely indecisive (panel F).

Like the preview effect, also the clear target orientation effect from Experiment 1 was replicated in Experiment 2. Responses in the tilt discrimination task were faster, *F*(1,18) = 14.23, *p* = .001, BF = 10.00, and clearly more accurate, *F*(1,18) = 36.94, *p* < .001, BF > 100, for upright than for inverted target faces. Also the fixation-locked EEG showed again a clear target face inversion effect from about 150 ms onwards that further extended across the whole post-fixation period. Importantly, the target orientation effect was present in the fN170 component consisting in a more negative deflection for inverted compared to upright target faces (BF evidence in Figure 9A, ERPs in Figure 9E). This effect was confirmed in an Anova at PO8, time window 165 to 250 ms, with *F*(1,18) = 14.54, *p* = .001.

Additionally, error rates indicated an interaction of Preview and Target Orientation factors, *F*(1,18) = 7.00, *p* = .016, which can be interpreted as a Preview Orientation main effect. This effect indicated slightly higher error rate with inverted (21.8%) compared to with upright (20.5%) preview faces. The BF for this effect was, however, indecisive and, if anything, suggested the absence an effect, BF = 0.47. We, therefore, do not consider this small effect (1.3% points difference) as very reliable.

As in Experiment 1, the early fixation-locked EEG also showed a clear Preview x Target Orientation interaction, equivalent to a Preview Orientation main effect, starting around 50 ms and extending to 170 ms post fixation onset (Figure 9C). As can be seen from Figure 9E, this effect indicated a more negative P1 with inverted compared to with upright preview faces, although the preview face was replaced by the target face at that point of the trial and the target face could have had a different overall orientation.

Again as in Experiment 1, evidence for the Preview x Target Orientation interaction became positive a second time around 350 ms at a set of central-parietal electrodes (Figure 9C). Again evaluated at electrodes CPz in the time window 320 to 400 ms, the target orientation effect consisting in a stronger negativity for inverted compared to upright targets, main effect *F*(1,18) = 5.59, *p* = .030, was more pronounced with an invalid (−1.20 μV) compared to with a valid preview (0.13 μV), *F*(1,18) = 11.49, *p* = .003, which likely again reflected increased processing of the target face orientation if the target presented new information different from the preview face. Thus, overall the results of Experiment 2 confirmed the main effects found in Experiment 1.

### 3.5. Experiment 2: The proportion manipulation affected tilt discrimination performance and the fixation-locked EEG, but it did not modulate the fN170 preview effect

Experiment 2 tested whether the preview effect found in Experiment 1 is the result of a more extensive prediction mechanism across trials, in the sense that it is influenced by expectations based on the frequency of events over an extended period of time rather than a single saccade. If the preview effect results from such a prediction mechanisms, then it should be larger in a block with mostly valid trials (66.6% valid, 33.3% invalid) than in a block with mostly invalid trials (33.3% valid, 66.6% invalid) (Figure 3). We, therefore, expected to find a Preview x Proportion interaction in the behavioral data of the tilt discrimination task and in the fN170 component of the fixation-locked EEG.

Interestingly, some hint for a Preview x Proportion interaction was provided by response times, *F*(1,18) = 5.64, *p* = .029, suggesting a slightly larger preview effect (57 ms) in the mostly valid block compared to the mostly invalid block (34 ms), which was the expected direction of the effect. However, the corresponding BF = 0.29 suggested no effect of this interaction, which renders the evidence rather uncertain. Another inconsistency in the response time data manifested in the main effect of Proportion which was not significant, *F*(1,18) = 2.14, *p* = .161, but exhibited a relatively high BF = 38.23.

In the error rates, the Preview x Proportion interaction was not significant, *F*(1,18) < 1, absence of effect confirmed by BF = 0.33, and also the Proportion main effect was not significant, *F*(1,18) = 0.05, *p* = .828, absence of effect confirmed by BF = 0.18.

In contrast to these equivocal behavioral results, the EEG data provided compelling evidence for the same fN170 preview effect in both mostly valid and mostly invalid blocks. BF values less than 1/3 at posterior lateral electrodes, where the fN170 preview effect is located, indicated the clear absence of a Preview x Proportion interaction (Figure 8F). This interaction was also not significant in a repeated measures Anova on average ERPs at PO8 from 165 to 250 ms, *F*(1,18) = 0.32, *p* = .581, at PO7, *F*(1,18) = 0.57, *p* = .462. As can be seen from the ERPs in Figure 8G, the difference in the amplitude between valid (solid line) and invalid trials (dashed line) was the same in mostly valid and in mostly invalid blocks. This crucial result suggests that the trans-saccadic preview effect in the fN170 component is not the result of context-sensitive predictions, which contrasts ideas about the predictive nature of the N170 (Johnston et al., 2017).

One might argue that the proportion manipulation was simply not strong enough to trigger a change in the fN170 preview effect. The proportion manipulation had, however, a pronounced influence on the fixation-locked EEG, in particular contralateral to the target face (right hemisphere) at posterior electrodes (Figure 9B). The direction of this effect at electrode PO8 is illustrated in Figure 8G. A more negative fN170 component occurred in the mostly valid than in the mostly invalid block, further corroborated by an Anova on average ERPs at PO8, time window 165 to 250 ms, *F*(1,18) = 12.77, *p* = .002. This clear influence of the proportion manipulation evidences that the 66.6% versus 33.3% manipulation was indeed strong enough to affect the fixation-locked EEG, showing that the proportion manipulation did influence face preview processing, but still it did not modulate the fN170 preview effect.

Apart from these Proportion effects of main interest, the factor Proportion interacted with Target Orientation later in the fixation-locked EEG and, surprisingly, in ipsilateral electrodes (Figure 9D, 9G). The effect was significant in an Anova on average ERPs at PO7, time window 550 to 800 ms, *F*(1,18) = 6.34, *p* = .021, suggesting that the late target face orientation effect was larger in the mostly valid than in the mostly invalid block. This effect probably indicates some variation in higher-level processing of the target face depending on the long-run frequency of valid and invalid trials. The reasons for its direction and for its ipsilateral location are, however, unclear. In any case, this finding does not influence our conclusions about the preview effect and its modulation by proportion.

### 3.6. Experiment 2: Evidence for pre-saccadic expectations in the preview-locked EEG response

If the proportion manipulation consisting in a block of mostly valid and a block of mostly invalid trials introduced expectations about the validity of a single trial, the preview face might have already been processed differently in mostly valid compared to mostly invalid blocks. Thus, expectation or prediction effects might already be present before the eye-movement during the preview period. We, therefore, analyzed the pre-saccadic period of the EEG signal, time-locked to the preview face display onset, with the factors Preview Orientation (upright, inverted), Proportion (mostly valid, mostly invalid), and also Target Orientation (valid, invalid). It is important to note that target orientation was unknown during the preview period and that the preview face was actually task-irrelevant since the task only involved the tilt of the post-saccadic target stimulus.

First, we found a classical N170 face inversion effect in response to preview face orientation as expected from an EEG study using face stimuli. Strong evidence from a whole-scalp BF (Figure 8A) demonstrated a more pronounced N170 for inverted compared to upright preview faces (Figure 8C). This effect was corroborated by an Anova on average ERPs at PO8, from 200 to 260 ms, *F*(1,18) = 29.63, *p* < .001. Compared to previous EEG studies on face perception showing an onset of the N170 largely around 150 to 200 ms (Bentin et al., 1996; Eimer, 2000; Eimer et al., 2010; Roxane J Itier and Taylor, 2004; Roxane J. Itier and Taylor, 2004; Rossion et al., 2000; Towler et al., 2012; Watanabe et al., 2003), our N170 appeared rather late at 200 ms (Figure 8A). This discrepancy might be explained by a difference in stimulus material. Previous studies on the N170 usually presented faces at the fovea in portrait format (for an exception see Pajani et al., 2017). The faces in our study were cut-outs excluding hair and the shape of the head, presented in the periphery, which might have slowed down face recognition processes and therefore might have led to a later N170 face inversion effect.

Instead of impacting on early stages of post-saccadic processing, the proportion manipulation influenced later stages of a face inversion effect. In about the second half of the preview period, an inverted preview face led to a more negative deflection than an upright preview face (Figure 8A, 8C), corroborated by an Anova on average ERPs at PO8, from 300 to 450 ms, *F*(1,18) = 21.70, *p* < .001. This effect possibly reflected a modulation of the N250 or N400 face processing components. Interestingly, as can be seen from Figure 8C, this late preview face orientation effect declined earlier in the mostly invalid than in the mostly valid block. In particular, between cue onset (at 500 ms) and saccade onset (see the histogram of saccade latencies in Figure 8D) the preview face orientation effect was gone in the mostly invalid block but still present in the mostly valid block. This earlier reduction of the preview face orientation effect in the mostly invalid compared to the mostly valid blocks around the time of cue onset is further illustrated in the scalp maps in Figure 10. BF evidence for the corresponding Preview Orientation x Proportion interaction is presented in Figure 8B. An Anova on average ERPs at PO8, 450 to 600 ms post preview onset, corroborated this interaction, *F*(1,18) = 16.99, *p* = .001. Critically, this effect could not simply be explained by a difference in saccade latencies between mostly valid and mostly invalid blocks, because saccade latencies did not differ between Preview Orientation and Proportion conditions: Proportion main effect, *F*(1,18) = 0.63, *p* = .439, BF = 1.14, Preview Orientation main effect, *F*(1,18) = 0.14, *p* = .714, BF = 0.17, Preview Orientation x Proportion, *F*(1,18) = 0.00, *p* = .997, BF = 0.24. As expected, also the factor Target Orientation did not affect saccade latencies, all *ps* > .089, all BFs < 0.29. The more sustained preview orientation effect in the mostly valid compared to the mostly invalid block might have, thus, reflected expectations about the upcoming target orientation based on the pre-saccadic input.

**Figure 10.**
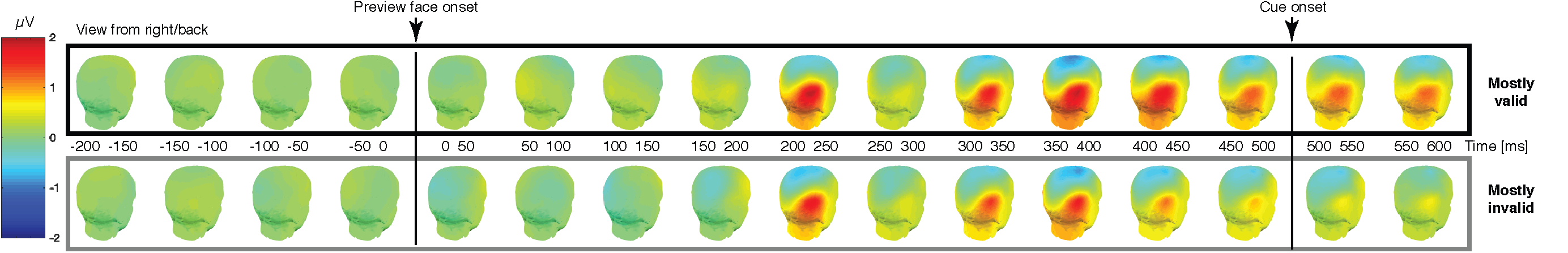
Scalp map of the preview-display-onset locked face inversion effect at lateral posterior sites (upright minus inverted). In the mostly valid block (upper row) the late face inversion effect remained, whereas it declined before cue onset and disappeared with cue onset in the mostly invalid block (lower row). Evidence for the corresponding Preview Orientation x Proportion interaction in Figure 8B.

Apart from these effects of main interest, the whole-scalp analysis of the pre-saccadic period revealed also a main effect of Proportion (Figure 11A), and some unsystematic effects involving Target Orientation (Figure 11B-E). The main effect of Proportion simply suggested a more positive ERP primarily at PO10 and at central-parietal electrodes in the mostly invalid compared to the mostly valid condition between cue onset and saccade onset, corroborated by an Anova on average ERPs, 500 to 650 ms after preview onset, at PO10, *F*(1,18) = 17.54, *p* = .001. This effect emphasizes that the influenced of Proportion on the EEG response in general. Compared to the other effects observed in this dataset, the effects involving Target Orientation were very short-lived and their spatiotemporal pattern varied considerably (Figure 11B-E).

**Figure 11.**
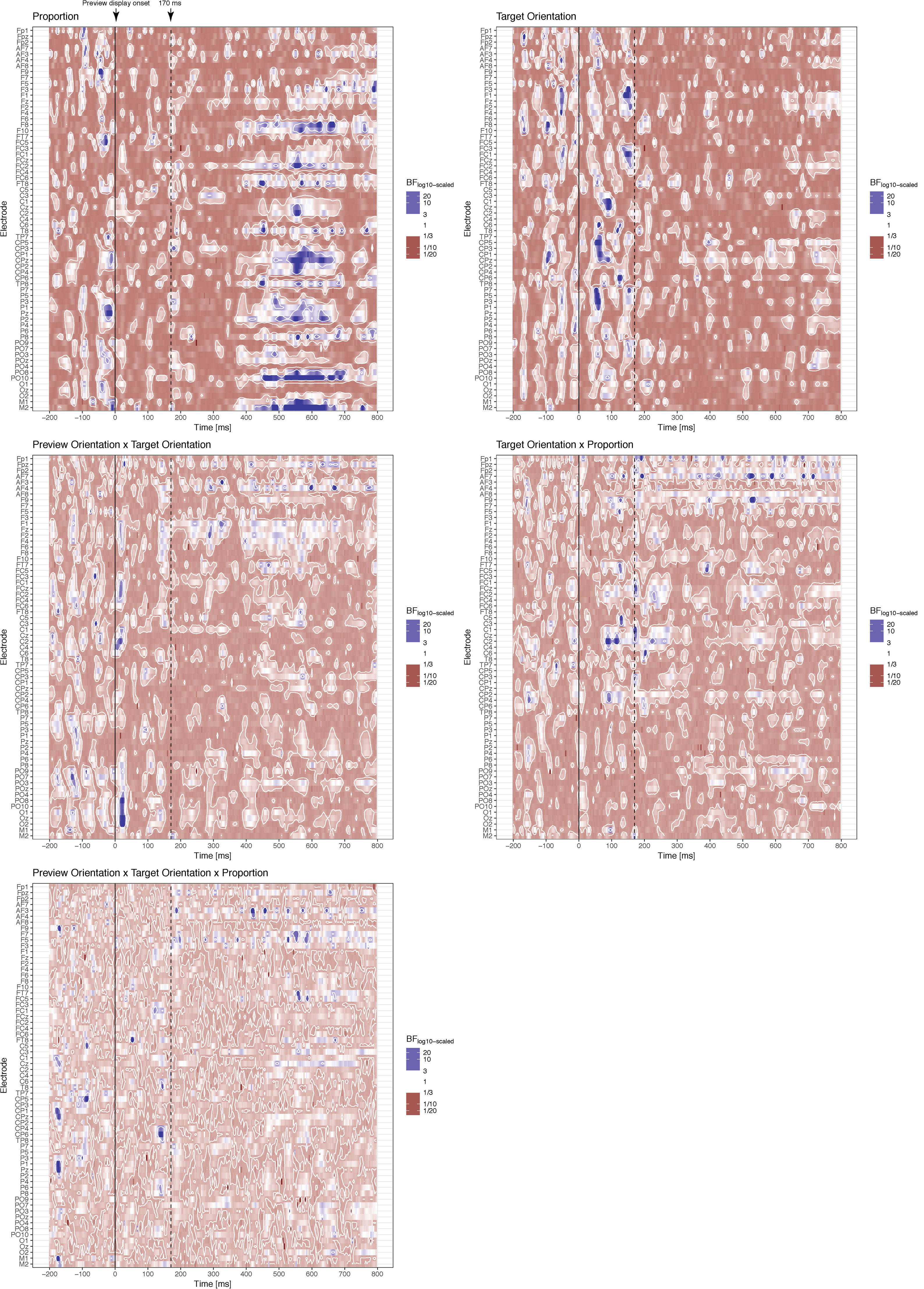
Preview-onset-locked whole-scalp Bayes factor (BF) for the remaining main and interaction effects of Experiment 2 not illustrated in Figure 8. Some positive evidence for a main effect of proportion was present primarily at PO10 and some central-parietal electrodes (panel A). The other effects involving Target Orientation (panel B-E) showed spatio-temporally extremely limited and unsystematic patterns of occasional positive evidence.

### 3.7. Experiment 2: The onset of the N170 face inversion effect in the preview period was later than the onset of the fixation-locked fN170 face inversion effect

As can be seen from Figure 8, the N170 elicited by the onset of the preview display appeared a bit later than the fixation-locked N170 (see in particular Figure 8C and 8G). To determine the statistical evidence for this effect, we computed onset latencies of the face inversion effect expressed as difference waveform between trials with upright and inverted faces at electrode PO8. We computed upright-minus-inverted preview orientation ERPs separately for mostly valid and mostly invalid blocks for the preview-display-locked data. For the fixation-locked data, we computed upright-minus-inverted target orientation ERPs separately for mostly valid and mostly invalid blocks and also separately for trials with valid and invalid preview. The design for the latency onset analysis was, thus, a 2 (Proportion: mostly valid, mostly invalid) by 3 (Preview: valid/fixation-locked, invalid/fixation-locked, none/preview-locked) design. Onset latencies of the face inversion effect were defined via a 50% peak amplitude criterion based on jack-knifed subsamples. In other words, the onset latency was the time stamp of the sample at which the leave-one-participant-out averaged difference waves between upright-minus-inverted face ERPs reached the value closest to 50% of its maximum activation within 100 to 250 ms after preview-display-onset/fixation-onset (Miller et al., 1998; Ulrich and Miller, 2001). These latency onset values were subjected to a repeated measures Anova with the factors Preview (valid, invalid, none) and Proportion (mostly valid, mostly invalid). The resulting F and p-values were corrected for the reduced error introduced by jack-knifing (Ulrich and Miller, 2001). It is at present unclear how a Bayes factor would have to be corrected for the reduced error due to jack-knifing. To avoid this issue, we applied the correction factor that counteracts the reduction in error, (n-1)^2 (Ulrich and Miller, 2001, see in particular Appendix), to the error sum of squares term obtained from the Anova, which further allowed Bayes factor approximations (Huber-Huber, 2016; Masson, 2011; Nathoo and Masson, 2016; Wagenmakers, 2007).

This latency onset analysis of the preview-locked and the fixation-locked face inversion difference waves showed a main effect of Preview (valid/fixation-locked, invalid/fixation-locked, none/preview-locked), *F*(2,36) = 27.18, *p* < .001, BF_approx_ > 100. Post-hoc tests based on Scheffe’s interval as critical difference (Ulrich and Miller, 2001) revealed a significant difference (at alpha-level .05) between the valid/fixation-locked and the invalid/fixation-locked face inversion effect, between the valid/fixation-locked face inversion effect and the face inversion effect without preview, but not between the invalid/fixation-locked face inversion effect and the face inversion effect without preview (Figure 12). Both the factor Proportion, *F*(1,18) = 0.70, *p* = .413, BF_approx_ = 0.330, and the Preview x Proportion interaction, *F*(2,36) = 0.15, *p* = .863, BF_approx_ = 0.031, were not significant.

**Figure 12.**
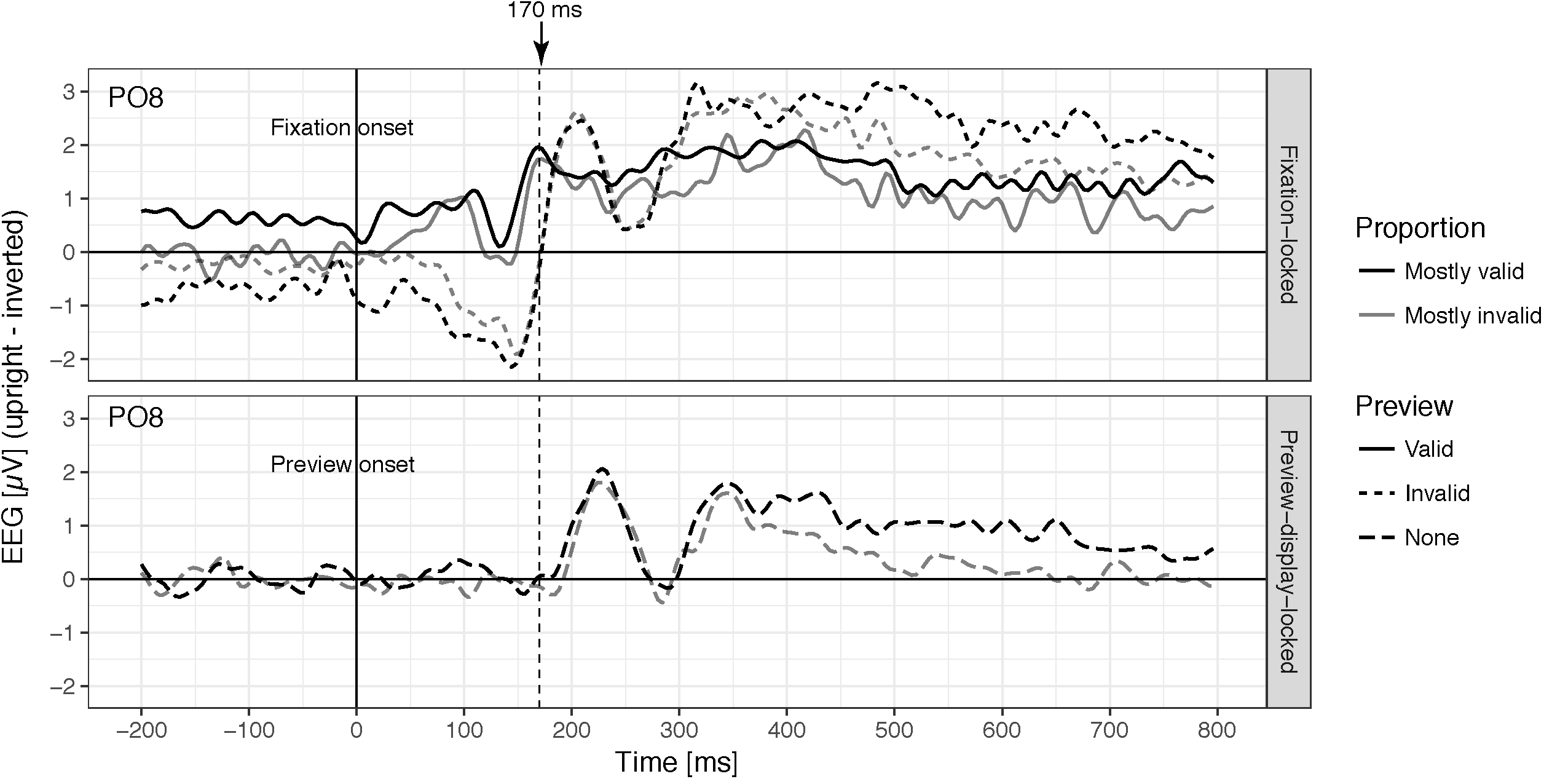
Time course of the face inversion effect calculated as difference between ERPs to upright faces minus ERPs to inverted faces separately for fixation-locked data (upper panel) and preview-display onset locked data averaged across both target face orientation (lower panel). The onset of the face inversion effect was earliest in the post-fixation period with a valid preview peaking at 170 ms (solid lines, upper panel). In contrast, an invalid preview delayed the face inversion effect (dashed lines upper panel). The latest face inversion effect occurred in response to the preview display, that is, before any eye movement was made (lower panel). The Proportion factor did not affect face inversion effect latency.

## 4. Discussion

We investigated the time course of trans-saccadic perception in a combined EEG and eye-tracking study. In Experiment 1, we established a preview effect in behavioral data and in the lateralized posterior fN170. Participants were more efficient in discriminating target-face tilt with a valid preview than with an invalid preview. In line with this result, the fN170 component was clearly more pronounced with an invalid than with a valid preview, which is the same effect direction as the preview positivity known from reading research (Dimigen et al., 2012, in particular Figure 3B). Our preview effect with faces emerged, however, much earlier than the preview positivity for reading (ca. 120 ms versus ca. 200 ms post fixation). We also found a later central-parietal effect similar to the later and more central preview component in reading research (Dimigen et al., 2012, Figure 3B). Again our late effect started earlier and instead of consisting in a Preview main effect it consisted in a Preview x Target Orientation interaction suggesting more in-depth processing of the target face orientation with invalid compared to with valid preview. These results suggest that trans-saccadic integration effects can be found at different temporal scales for different types of stimuli, possibly related to the different timing for processing these stimuli (Herrmann et al., 2005; e.g. Sereno and Rayner, 2003).

In addition to the trans-saccadic preview effect in the fN170, we found a clear face inversion effect (Bentin et al., 1996; Eimer, 2000; Eimer et al., 2010; Roxane J Itier and Taylor, 2004; Roxane J. Itier and Taylor, 2004; Rossion et al., 2000; Towler et al., 2012; Watanabe et al., 2003). This effect was also clearly present in response times and error rates in the expected direction of better performance with upright than with inverted target faces. Importantly, the target orientation and preview effects were additive suggesting that they reflect two independent processing stages, one for face detection and one for trans-saccadic integration. Only at a later stage, target orientation and preview interacted, which could mean that the outcomes of the two separate early processes are combined at this later stage: if there was a change during the saccade, then target face orientation received more in-depth processing.

In addition to increasing the amplitude of the fN170 in general, an invalid preview also delayed the face inversion effect to a similar onset as the face inversion effect triggered by the preview face itself. This result suggests that EEG studies in controlled experimental settings without eye movements underestimate the latency of visual EEG components in normal viewing, because real-world perception is usually preceded by a pre-saccadic preview, resembling the valid condition. In contrast, most experimental settings that prevent eye movements are like the preview onset locked condition, which triggered a later N170 face inversion effect compared to the fixated-evoked effect.

In Experiment 2, we asked whether the beneficial effect of the preview for post-saccadic processing, in particular on the fN170 component, was the result of a contextually-sensitive prediction process. In other words, does the trans-saccadic effect across a single eye movement take into account the frequency of valid and invalid trials? The direction of the fN170 preview effect, with a larger fN170 for invalid than for valid conditions, is consistent with a prediction error signal (Friston, 2010, 2005; Friston et al., 2012; Summerfield and Egner, 2009). If the fN170 preview effect reflected a context-sensitive predictive process, we reasoned that it should adapt to the frequency of events such that it would become larger in a block with more valid trials and smaller in a block with more invalid trials (Summerfield et al., 2008). The results of Experiment 2, however, contradicted this idea: The same preview effect was found in both blocks and confirmed by strong statistical evidence from a Bayes factor analysis. Our results, therefore, indicate that the fN170 preview effect occurs regardless of context or recent experience, making it different from many classical prediction effects. Still, we observed a more negative fN170 in the mostly-valid compared to mostly-invalid block, which suggests that the proportion manipulation with 33.3% versus 66.6% was strong enough to be picked-up by the participants and influence face processing to some extent, though not impacting the preview effect.

In the response time data, the effect of the proportion manipulation was less clear. The Anova provided some hint for a larger preview effect with mostly valid than with mostly invalid trials. The BF, however, provided negative evidence casting some doubt on the Anova result. In error rates, there was clearly no such modulation. This discrepancy between the behavioral and the EEG data suggests that behavior in the task was not only determined by the early stages of post-saccadic processing reflected in the fN170. It is useful to note, in this context, that the preview in our design was task-irrelevant, since the response was based on information (tilt of the target face) which was only present in the target and not in the preview.

Our results contrast previous notions of the N170 as being related to visual predictions. In an elegant study, Johnston and colleagues (2017) showed that violating visual predictions derived from a sequences of image changes viewed without eye movements elicited an N170. They even proposed to use this component to study sensory predictions across saccadic eye-movements. Moreover, the source of visual prediction errors signals has been localized in the fusiform face area (de Gardelle et al., 2013a, 2013b) which has also been identified as one of the neural generators of the N170 component (e.g. Corrigan et al., 2009). Our results, however, necessitate a reconsideration of the function of the N170/fN170 in predictive perception, in the classical sense of predictions that take account of context and recent experience.

One possibility is that predictions across saccadic eye movements (Edwards et al., 2017; Ehinger et al., 2015) might not obey the same principles as concurrent sensory predictions in the visual system without saccades (Alink et al., 2010; Johnston et al., 2017). This conjecture implies that the N170 and the fN170 respond differently to the same type of prediction manipulation, which has not yet been tested.

An alternative is that, although all types of prediction and expectation effects are based on the regularities and statistics of the environment, there are numerous ways of how these effects can be introduced (De Lange et al., 2018) and this might have implications for the precise neural mechanism that is targeted by the prediction manipulation. Johnston and colleagues (2017) studied visual prediction error signals by contrasting predictable and unpredictable image transitions within systematic sequences of images. The frequency of predictable and unpredictable trials was, however, balanced. In the present study, we manipulated the frequency of valid and invalid trials. This methodological difference could have been critical for the discrepant findings.

Finally, although proportion manipulations of 25% versus 75% have been successful in the past (Summerfield et al., 2008) and our proportion manipulation with 33.3% versus 66.6% was similar, it might still not have been strong enough to trigger an adaptation of trans-saccadic predictions (Kovács and Vogels, 2014; Mayrhauser et al., 2014). It is well-known that effects of expectation scale with validity of the prediction just like endogenous attention scales with cue validity (Giordano et al., 2009; Kok et al., 2012). Hence, more extensive training with trans-saccadic changes than the one realized in the present design (e.g. Herwig et al., 2015; Valsecchi and Gegenfurtner, 2016) might modulate the fN170 preview effect.

Overall, our results are consistent with the idea of two stages of visual predictions. The trans-saccadic preview effect found relatively early (100 – 170 ms) was independent of the proportion manipulation. It might be relatively automatic and resistant to change over a brief time period of a few trials. In terms of the second stage, the preview face inversion effect before the saccade was more sustained in blocks with mostly valid compared to blocks with mostly invalid trials. This result suggests that, based on the proportion manipulation, there was an expectation for the same face orientation again as target in the mostly valid block, less so in the mostly invalid block. Because saccades are executed in sequence in natural vision, a pre-saccadic effect could be considered as a late post-saccadic effect. With this assumption, our findings are consistent with the idea that later stages of perceptual processing are more susceptible to global stimulus regularities than early stages like the one of the fN170 (see also Pajani et al., 2017; Summerfield et al., 2011).

In any case, the preview effect in the fN170 can still be interpreted as a prediction error in terms of predictive coding (Grotheer and Kovács, 2016). In a computational sense, predictive coding only means that, instead of transmitting the complete bottom-up signal from lower to higher processing levels, only the prediction error is propagated in a feed-forward fashion (Friston, 2010; Spratling, 2017). Predictive coding does not imply anything about critical frequencies of events required for adjusting top-down predictions. Thus, even though the proportion manipulation did not influence the fN170 preview effect, the preview effect itself might still have resulted from predictive coding circuits (Bastos et al., 2012), with these circuits not influenced by our proportion manipulation.

In conclusion, the current results show a strong effect of a task-irrelevant preview face on post-saccadic face processing. We make about three saccades every second, and it takes about 100 ms until visual information arrives at object recognition areas (Foxe and Simpson, 2002). If there was no perception during that time we would be blind for about four hours each day (Melcher and Colby, 2008). Our results confirm that perception does not start anew with a new fixation. What we see in the periphery before we make an eye-movement affects post-saccadic processing. Moreover, our data even showed a *preview face* orientation effect in the early stage of *post-saccadic* processing (cf. Mirpour and Bisley, 2016). This particular result suggests that, instead of being blind after fixation onset, we perceive what was there before the eye movement which, in natural viewing, is also what will be there at the beginning of the new fixation. However, only after roughly 100-120 ms post-saccadic visual processing reflects what is actually in front of our eyes.

## Acknowledgements

The research was supported by a European Research Council (ERC) grant, “Construction of Perceptual Space-Time” (StG agreement no. 313658) awarded to David Melcher. Many thanks to Matteo Valsecchi and Alexander Schütz for insightful discussions, and to Marco Coratolo, Xenia Dmitrieva, and Gianmarco Maldarelli for help with data collection.

## Footnotes

1 We checked the equivalence of the Preview Orientation main effect and the Preview x Target Orientation interaction explicitly with two Anovas computed on the average amplitude within 300-400 ms post fixation onset at electrode pair PO7/8. One Anova contained the effect of Preview Orientation whereas the other Anova coded the same data with the effect of Preview instead. The first Anova showed a main effect of Preview Orientation with the values *F*(1,17) = 4.39, *p* = .051. The second Anova showed a Preview x Target Orientation interaction with exactly the same values *F*(1,17) = 4.39, *p* = .051. Besides that, the main effect of Target Orientation was also exactly the same for both Anovas, *F*(1,17) = 8.92, *p* = .008. Clearly, the Preview Orientation main effect translates into a Preview x Target Orientation interaction, and vice versa.

